# Structure-guided molecular grafting of a complex broadly neutralizing viral epitope

**DOI:** 10.1101/860825

**Authors:** Goran Bajic, Max J. Maron, Ming Tian, Garnett Kelsoe, Masayuki Kuraoka, Aaron G. Schmidt

## Abstract

Antigenic variation and viral evolution have thwarted traditional influenza vaccination strategies. The broad protection afforded by a “universal” influenza vaccine will come from immunogens that elicit humoral immune responses targeting conserved epitopes on the viral hemagglutinin (HA), such as the receptor-binding site (RBS). Here, we engineered candidate immunogens that use non-circulating, avian influenza HAs as molecular scaffolds to present the broadly neutralizing RBS epitope from historical, circulating H1 influenzas. These “resurfaced” HAs (rsHAs) remove epitopes potentially targeted by strain-specific responses in immune-experienced individuals. Through structure-guided optimization we improved two antigenically different scaffolds to bind a diverse panel of pan-H1 and H1/H3 cross-reactive bnAbs with high affinity. Subsequent serological analyses from murine prime-boost immunizations show that the rsHAs are both immunogenic and can enrich for RBS-directed antibodies. Our structure-guided, RBS grafting approach provides candidate immunogens for selectively presenting a conserved viral epitope.

## MAIN TEXT

Influenza evolves primarily at the human population level and within its animal reservoirs (swine and avian)^1^. Host humoral pressure, which predominantly targets the viral hemagglutinin (HA), selects for influenza mutations that render previous immune responses suboptimal. The humoral response then evolves, through immune memory and further B cell affinity maturation ^2-5^. The net effect of this on-going selection across the entire population exposed to the virus is a virus-immunity “arms race”. The repeated exposure to influenza in the human population results in preexisting immunity which influences subsequent immune responses ^6-11^. This immunological memory ^12,13^ presents a significant hurdle towards the development of a “universal” influenza vaccine. Strategies are necessary that both overcome the recall of refined, strain-specific responses and elicit broadly neutralizing antibodies (bnAbs).

bnAbs against influenza HA target two relatively invariant epitopes the receptor binding site (RBS) on the HA “head” and a surface along the HA “stem” ^14^. While stem-directed immunogens are in clinical development, efforts focusing on the RBS have lagged behind ^14^. A significant challenge for RBS-directed immunogens is presentation of the complex RBS structure that includes multiple segments, separated in linear sequence, but adjacent in conformational space ^15^. While computational design of novel protein scaffolds has been done for HIV and RSV, the grafted epitopes were often less-complex (e.g., a single alpha-helix) ^16,17^.

To overcome the significant hurdle of *de novo* protein design we hypothesized that the RBS epitope from one HA subtype could be transplantable onto another antigenically distinct HA. We used non-circulating, avian influenza HAs as molecular scaffolds to present the RBS from circulating H1 influenzas. These resurfaced HA (rsHA) scaffolds present the H1 conserved RBS recognized by bnAbs and remove other epitopes targeted by strain-specific responses in immune-experienced individuals. The crystal structure of one scaffold in complex with a bnAb allowed for further structure-guided optimization of two antigenically distinct scaffolds to bind a diverse panel of pan-H1 and H1/H3 cross-reactive bnAbs. Immunization with a recombinant H1 HA followed by a single, heterologous boost with our rsHA immunogen showed comparable levels of RBS-directed antibody response to the H1 homologous prime-boost regimen. These data suggest that these rsHA immunogens with further optimization of the vaccine regimen may provide a pathway to a universal influenza vaccine, by exploiting the immunogenicity of the conserved RBS.

## RESULTS

### Grafting the H1 SI-06 RBS onto acceptor HA scaffolds

As a proof of principle, we chose the circulating H1 RBS epitope as the basis of a “donor” graft to scaffold onto HA subtypes not currently circulating in the human population (**Fig. 1a**). H1 influenzas can be grouped into roughly three antigenic “clusters” with prototypical members represented by H1 Massachusetts/1/1990 (H1 MA-90), H1 Solomon Islands/03/06 (H1 SI-06) and H1 California/04/2009 (H1 CA-09) (**Fig. 1b** and **Extended Data Fig. 1**) ^6^. Importantly, bnAb have been identified that can span these antigenic clusters (e.g., CH67 ^18^, 641 I-9 ^19^, Ab6639 ^20^ and 5J8 ^21^). We used the H1 SI-06 as the initial donor and defined four “segments”, S1-S4, comprising the RBS epitope for grafting (**Fig. 1b**). These segments include 7 of the 13 critical residues that contact the receptor, sialic acid; these 13 residues define the RBS “core” (**Extended Data Fig. 1**). Many of the remaining residues not included in the graft (e.g., Y95 and W153) are in the base of the RBS and are nearly invariant across influenza subtypes. For the initial molecular scaffolds, we selected two non-circulating group 2 influenzas H4N6 A/America black duck/New Brunswick/00464/2010 (H4 NB-10) and H14N6 A/mallard/Wisconsin/10OS3941/2010 (H14 WI-10). We selected them because have little sequence similarity to circulating H1s (**Extended Data Fig. 2**). The acceptor HA S1-S4 boundaries were defined by aligning the H1 SI-06 sequence (**Fig. 1c**). The rsHAs have the following nomenclature: “rsH4NBvX”; the resurfaced (rs) HA scaffold subtype (H4), with an abbreviated strain name (NB) and different versions (*v*X). We could successfully overexpress the inter-group transfer of the H1 SI-06 RBS graft onto the H4 and H14 scaffolds resulting in our first generation rsH4NBv1 and rsH14WIv1 scaffolds (**Extended Data Fig. 3**).

**Figure 1:**
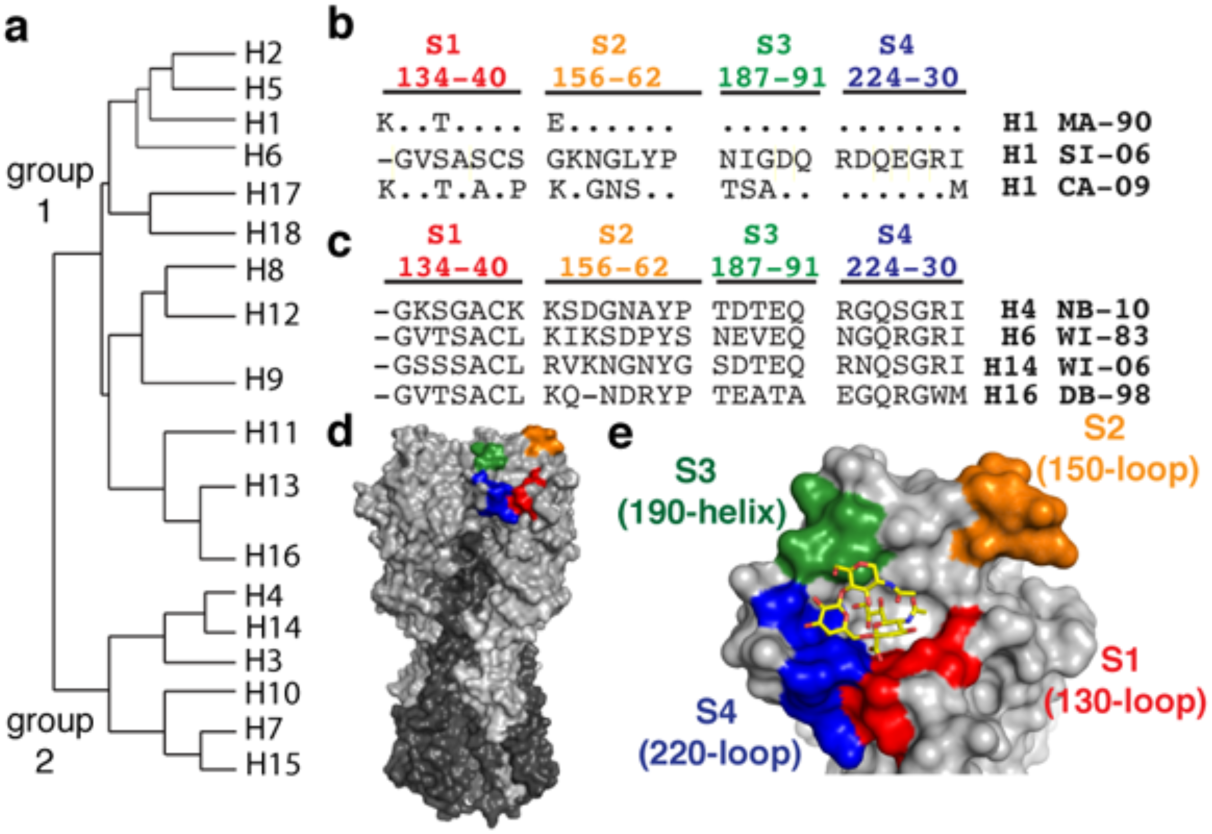
RBS grafts and sequence alignments. **a**, Phylogeny of influenza subtypes. Group 1 and 2 influenzas are annotated. **b**, Representative H1 antigenic clusters: H1 Massachusetts/1/1990 (MA-90), H1 Solomon Islands/03/2006 (SI-06) and H1 California/07/2009 (CA-09) are listed. Sequence alignment is in reference to SI-06 and conserved residues are marked as (.); segments defining the H1 RBS graft, S1-4, are colored. **c**, Residues comprising S1-4 of the acceptor scaffolds from non-circulating influenzas H4 New Brunswick/00464/2010 (H4 NB-10), H6 Wisconsin/617/1983 (H6 WI-83), H14 Wisconsin/10OS3941/2006 (H14 WI-06) and H16 Delaware Bay/296/1998 (H16 DB-98). **d**, Influenza HA trimer (PDB 5UGY) in surface representation. HA1 is in silver, HA2 in dark gray and S1-4 are colored. **e**, LSTc (stick representation) modeled in complex with HA. S1-4 is colored and HA is in silver.

### Binding affinities of bnAbs to rsHAs

We determined the binding affinities of a panel of RBS-directed Fabs to these initial scaffolds using by biolayer interferometry (BLI). This panel included four pan-H1 (CH67, 641 I-9, H2526 and H2227) and two H1/H3 cross-reactive bnAbs (K03.12 and C05) that engage the RBS ^19,21-24^. Each antibody has a footprint that overlaps with the RBS core but has different angles of approach and peripheral contacts (**Extended Data Fig. 4**). As seen in **Table 1**, neither scaffold bound all the RBS-directed Fabs. rsH4NBv1 bound only K03.12 with an equilibrium dissociation constant (K_D_) ∼5.2x greater than wildtype H1 SI-06. rsH14WIv1 bound CH67, K03.12 and C05 with K_D_s ∼17x, ∼2.7x and ∼1.3x greater than wildtype H1 SI-06. None of the antibodies bound wildtype H4 NB-10 or H14 WI-10. These data suggest that there are peripheral residues in the first-generation scaffolds that impede bnAb binding and/or that the conformation of the RBS graft is being presented in an altered conformation.

**Table 1.** Affinity measurements of rsHAs to a panel of RBS-directed Abs. K_D_s (in *µ*M) were obtained by applying a 1:1 binding isotherm using vendor-supplied software with at least three independent concentrations. The V_H_ gene usages are listed for each antibody. K_D_s beyond the limit of detection are reported as >100 *µ*M. Values are for monomeric HA heads and Fabs.

|  | **CH67**<br>*V<sub>H</sub>1~2* | **641 I-9**<br>*V<sub>H</sub>4~59* | **H2526**<br>*V<sub>H</sub>1~69* | **H2227**<br>*V<sub>H</sub>4~4* | **K03.12**<br>*V<sub>H</sub>1~2* | **C05**<br>*V<sub>H</sub>3~23* |
| --- | --- | --- | --- | --- | --- | --- |
| **H1 SI-06** | 0.57 | 0.67 | 0.56 | 0.39 | 0.83 | 1.1 |
| **H4 NB-10** | >100 | >100 | >100 | >100 | >100 | >100 |
| **H14 WI-10** | >100 | >100 | >100 | >100 | >100 | >100 |
| **rsH4NBv1** | >100 | >100 | >100 | >100 | 4.3 | >100 |
| **rsH4NBv3** | 1.6 | 17.7 | 0.93 | 0.67 | 0.65 | 0.34 |
| **rsH14Wlv1** | 9.5 | >100 | >100 | >100 | 2.2 | 1.4 |
| **rsH14Wlv2** | 0.31 | 2.7 | 0.20 | 1.0 | 1.1 | 0.065 |

### Structure of rsH4NBv1 in complex with bnAb, K03.12

To identify further modifications on the first-generation scaffold that can be engineered to improve affinity to the bnAb panel, we determined the crystal structure of the rsH4NBv1 head in complex with the cross-reactive H1/H3 K03.12 antibody (**Fig. 2a**). Like the previously characterized C05 antibody, K03.12 engages the RBS with almost exclusively CDR H3–dependent contacts ^22^. The antibody contacts 14 of the 26 residues in the S1-S4 RBS graft. Additional contacts are made with conserved residues critical for sialic acid interactions in the base of the RBS including Y95, W153, T155 and H183. Comparison of the K03.12-rsH4NBv1 structure and the K03.12-H3 Texas/50/2012 complex shows a nearly identical approach with a slight twist and rocking of the V_H_-V_L_ towards HA about the principle axis (**Extended Data Figs. 5a, b**) ^23^. The contacting residues within the antigen-combining sites between the two structures are nearly identical (**Extended Data Figs. 5c, d**). A comparison of the rsH4NBv1 HA to a wildtype H4 A/duck/Czechoslovakia/1956 (PDB 5XL3) and wildtype H1 SI-06 (PDB 5UGY) shows a displacement about S1 (150-loop) with a shift of ∼3Å, however the overall conformation in the other grafted RBS segments is similar to that of the wildtype H1 SI-06 RBS (**Fig. 2c**).

**Figure 2:**
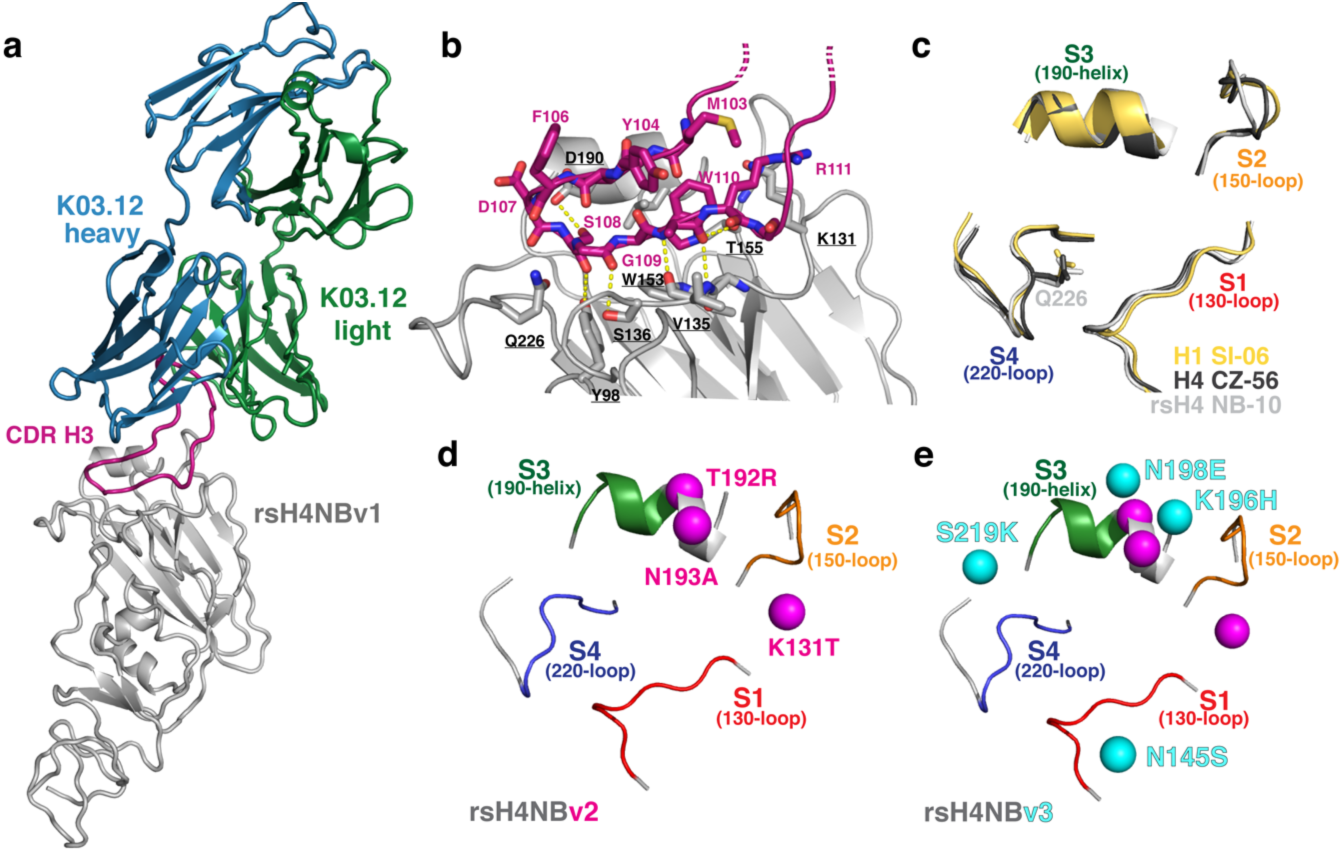
Structure of K03.12 in complex with rsH4NBv1 and scaffold improvement. **a**, Antibody K03.12 Fab (heavy and light chains are colored blue and green, respectively) in complex with rsH4NBv1 HA1 “head” (silver). The CDR H3 (magenta) is marked. **b**, Close-up of the antigen combining site. The CDR H3 (magenta) is shown in sticks with key interacting HA residues (silver). Hydrogen bonds are denoted in yellow, dashed-lines. **c**, Comparison of the RBS of H1 SI-06, yellow, (PDB 4YJZ), H4 A/duck/Czechoslovakia/1956 (H4 CZ-56), black (PDB 5XL3) and rsH4NBv1, silver. The segments of the grafts are labeled and colored. **d**, rsH4NBv2 and **e**, rsH4NBv3 with residues changed for each construct represented by a colored sphere. The original, unchanged segments from rsH4NBv1 are labeled and colored.

### Structure-guided improvement of the rsHA scaffolds

To improve scaffold binding to the RBS-directed bnAb panel, we docked the previously determined Fab structures onto the determined rsH4NBv1 structure to identify residues that may be modified to either alleviate steric clashes and/or reinforce interactions. We engineered two additional (scaffold) versions: 1) rsH4NBv2 had three mutations K131T, T192R and N193A (**Fig. 2d**) and 2) rsH4NBv3, had four additional mutations, N145S, K196H, N198E and S219K (**Fig. 2e**); no changes were made in the original grafted segments. We first assayed for scaffold improvement using an enzyme-linked immunosorbent assay (ELISA) (**Fig. 3**). None of the antibodies bound to wildtype H4 NB-10 (**Fig. 3a**) consistent with our BLI data using Fabs (**Table 1**). The second-generation scaffold increased affinity to four of the five Fabs (**Fig. 3b, c**) while the third generation, rsH4NBv3, resulted in high-affinity binding to all five RBS-directed antibodies (**Fig. 3c**). Based on this optimized rsH4NBv3 construct, we asked whether the same seven mutations could be made in context of the rsH14WIv1 scaffold to increase its affinity for the entire panel of RBS-directed antibodies. Indeed, when these mutations were engineered into rsH14WIv1 this optimized construct bound the entire panel of RBS-directed antibodies with high affinity (**Fig. 3g-i**). For reference, H1 SI-06 reactivity to the RBS-directed Ab panel is shown (**Fig. 3i**). A summary of the K_D_s are shown in **Figs. 3e, 3j**.

**Figure 3:**
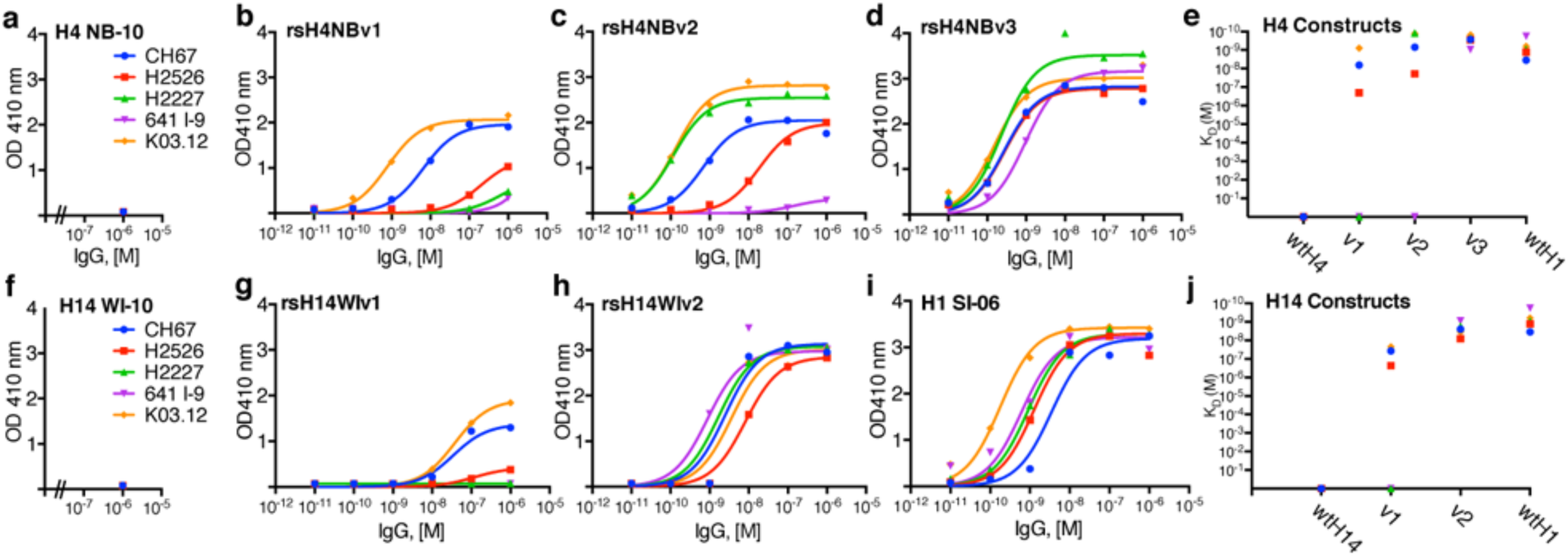
Reactivity of RBS-directed IgGs for rsHAs. CH67 (blue), H2526 (red), H2227 (green), 641 I-9 (violet) and K03.12 (orange) IgGs were titrated against **a**, wildtype H4 NB-10, **b**, rsH4NBv1, **c**, rsH4NBv1, **d**, rsH4NBv3, **e**, K_D_s obtained from curves in **a-d** for H4 constructs. IgG titrations for **f**, wildtype H14 WI-10, **g**, rsH14v1, **h**, rsH14v2 and **i**, control H1 SI-06 HA constructs. **j**, K_D_s obtained from curves in **f-i** for H14 constructs. ELISA measurements were done in duplicate over the concentration range except for **a**, and **f**, where only 1µM final concentration of IgG was tested. The wildtype H1 SI-06 values in **e** and **j** are both derived from **i**. Curve fitting and K_D_s were obtained using GraphPad Prism version 6.0.

Finally, we tested the optimized scaffolds in BLI to obtain more accurate K_D_s, free of avidity affects present when using IgGs in ELISA. As seen in **Table 1**, both optimized scaffolds significantly increased affinities over their first-generation counterparts; rsHAs based on both optimized scaffolds bound all six bnAbs included in the panel. In particular, the rsH14WIv2 scaffold yielded affinities for CH67, H2526 and C05 even greater than those of wildtype H1 SI-06. In some cases (e.g., C05) the optimized resurfaced scaffold bound the Fab >10-fold more tightly than did wildtype H1 SI-06. In general, the rsH14WIv2 had affinities closest to the wildtype H1 SI-06. These data suggest a set of key residues, in addition to the initial RBS-donor grafts, that can be grafted onto other potential scaffolds to present an “optimized” epitope to bind (or elicit) a diverse set of RBS-directed bnAbs.

### Serum and germinal center B cell responses to the rsHA immunogen

To evaluate whether our rsHA immunogens can enrich for RBS-directed responses by allowing the expansion and differentiation of memory B cells, we first generated a knock-in (KI) mouse that has a human J_H_6 segment; the human J_H_6 segment was chosen to mimic two key features of observed human-RBS directed bnAbs that is absent in the wildtype B6 murine model: 1) a poly-tyrosine motif and 2) an overall length of 19-20 amino acids necessary for engaging the RBS^19^. We therefore immunized these generated mice (see Methods) with recombinant H1 SI-06 HA, to prime humoral immunity to an H1, then boosted with our recombinant rsH4NBv3 (**Fig. 4a)**. We compared the resulting serum and GC responses to those originating from a homologous H1 SI-06/ H1 SI-06 or a heterologous H1 SI-06/rsH4NBv3 prime/boost immunization. We found that post-boost, the serum IgG titers, as well as the relative serum reactivity to the initial H1 SI-06 HA were comparable between the two cohorts (**Fig. S6a**). To deconvolute the contribution of the H4 scaffold-dependent Abs, we performed a serum competition assay with H1 RBS-directed bnAb CH67 (**Fig. 4b**) and found no demonstrable difference between the two cohorts. At the cellular level, we noted that the GC B cell frequencies were ∼3-fold higher in mice boosted with recombinant rsH4NBv3 (**Fig. 4c and S7**). These data potentially point to disparities in immunogenicity between H1 SI-06 and rsH4NBv3 and highlight the apparent immunodominance of the H4 scaffold. To quantify the serum abundance of RBS-directed antibodies, we screened the sera from the two cohorts against H1 SI-06 and rsH4NBv3 HAs in an ELISA (**Fig. 4d, e**). Sera from the heterologous immunization reacted more strongly with rsH4NBv3 (**Fig. 4e and S6b**) and as strongly as those from the homologous one with H1 SI-06 (**Fig. 4d and S6c**), indicating that the rsH4NBv3 boost elicited amounts of RBS-directed antibodies comparable to homologous boost, if not higher. These data highlight the potential benefit of boosting with rsH4NBv3 as means to direct antibody responses to the RBS.

**Figure 4:**
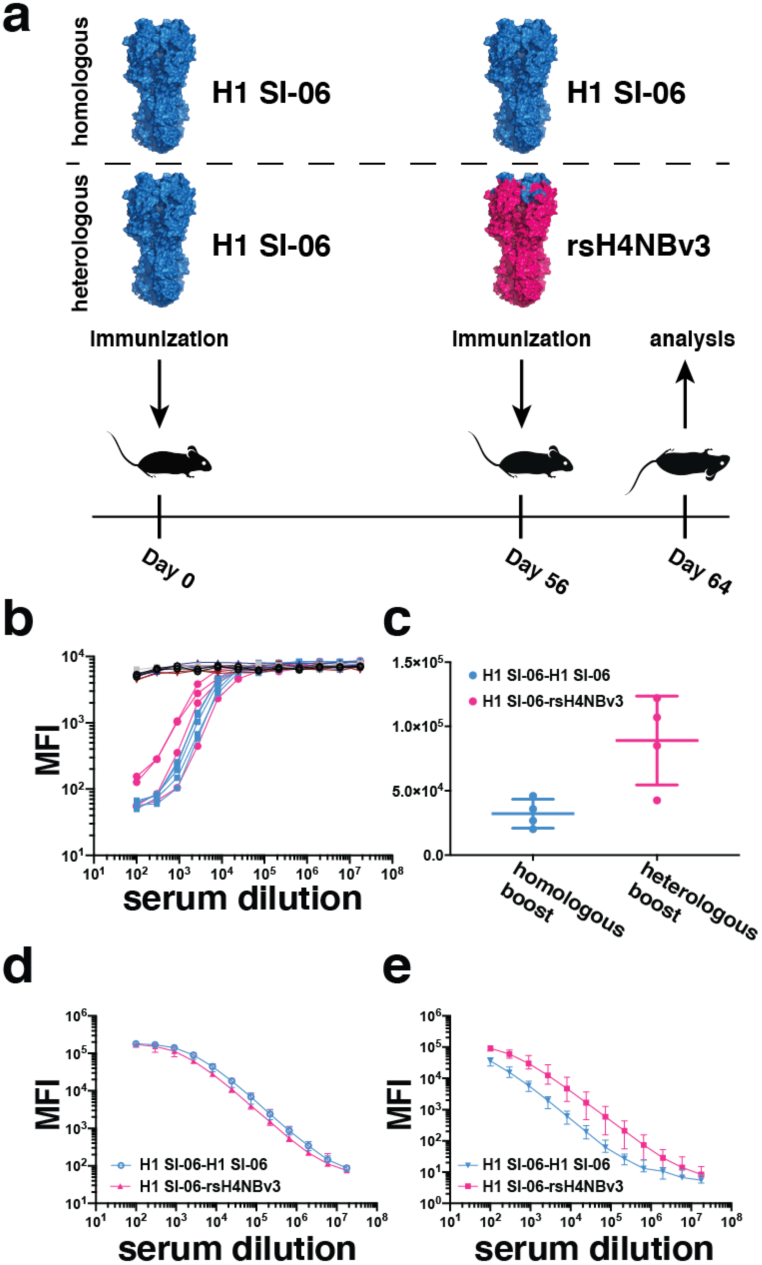
*in vivo* assessment of rsHA immunogenicity. **a**, schematic representation of the immunization strategy **b**, CH67 competition experiment against the sera from wild-type non-immunized B6 mice (black), J_H_6 mice primed and boosted with H1 SI-06 (blue) and mice primed with H1 SI-06, then boosted with rsH4NBv3 (pink), **c**, total GC B cell counts in the animal cohorts, **d**, serum reactivity against H1 SI-06, **e**, serum reactivity against rsH4NBv3.

## DISCUSSION

Current influenza research has focused on the development of a universal influenza vaccine. Such a vaccine should induce broad immunity a) within seasonal, circulating H1 and H3 subtypes, b) across subtypes (heterosubtypic) and c) pre-pandemic (e.g., H5, H7). The pathway to achieving this broad protection will likely come from eliciting or boosting humoral responses to conserved sites on HA such as the RBS and “stem”. Immunogen design strategies thus far have focused almost exclusively on targeting the conserved stem through either selectively displaying the HA2 stem ^25^ or using chimeric HAs that present circulating H1, H3 stems with a heterologous HA1 “head” ^14^.

The data presented here show an alternative strategy for candidate immunogens to focus the immune response to the broadly neutralizing epitope of the RBS. Grafting of the H1 SI-06 RBS epitope onto two antigenically distinct HA scaffolds exploits the overall architecture of the HA protein circumventing the significant challenge in *de novo* protein scaffold design. Through structure-guided engineering, the optimized scaffolds bind a diverse panel of pan-H1 and H1/H3 cross-reactive RBS antibodies that represent the type of response one might wish to elicit by a universal influenza vaccine. Collectively, the bnAbs in the panel (**Table 1**) bind all H1 isolates both pre-pandemic (<2009) and post-pandemic (>2009) as well as circulating H3 influenzas. Importantly, this collection of RBS-directed bnAbs tolerate the heterologous peripheries of the scaffolds surrounding the graft. These immunogens therefore would stimulate affinity maturation to refine humoral responses to the conserved, RBS core contacts shared between the scaffolds while adapting and accommodating antigenically distinct peripheries.

Our *in vivo* mouse immunization studies compared the resulting serum and GC responses from mice primed and boosted with either homologous or heterologous HA immunogens. Despite no discernable difference in a competition assay with a standard RBS-directed mAb CH67, we noted that the heterologous immunization elicited more robust GC B cell responses, owing either to the influx of new, H4-specific clones and/or recall of the H1 RBS-specific memory B cells. Although further studies involving fate labeling antigen-experienced cells and single-cell sequencing will be necessary to deconvolute these effects, we did note that the sera from the heterologous immunization reacted more strongly with the boosting antigen and as strongly as those from the homologous one with the priming antigen. These data indicate that the heterologous boost elicited comparable, if not higher, amounts of RBS-directed antibodies and highlight the benefit of boosting with rsH4NBv3 to direct antibody responses to the RBS.

A significant hurdle for the development of a universal influenza vaccine is preexisting immunity present in the human population either through repeated seasonal vaccination or influenza infection. The HA scaffolds derive from avian influenzas that, to date, have not circulated in the human population and thus would likely avoid boosting strain-specific, memory recall responses in immune-experienced individuals. The strategy described here could be used to direct naïve immune responses to the conserved RBS through a prime-boost vaccine approach and/or could boost subdominant RBS-directed bnAbs already present in immune-experienced individuals^11,19^. More generally, the immunogen design approach of epitope grafting could be used for other rapidly evolving pathogens for which preexisting immunity is present (e.g., RSV and dengue).

## METHODS

### Expression and purification of HA

rHA1 and “head” and rHA full length soluble ectodomains (FLsE) constructs were cloned into pFastBac vector for insect cell expression (Hi5 cells) or pVRC vector for mammalian expression (293F or 293T cells). HAs were derived from the following templates: H4N6 A/America black duck/New Brunswick/00464/2010 (GenBank: AGG81749.1) and H14N6 A/mallard/Wisconsin/10OS3941/2010 (GenBank: AGE03043). All constructs were confirmed by DNA sequencing at the DNA Sequencing Core Facility at Dana Farber Cancer Institute. For biolayer interferometry (BLI) and crystallography the HA1 head constructs contained a HRV 3C-cleavable C-terminal His_6X_ tag or SBP-His_8X_tag. The HA FLsE constructs used in ELISA assays contained a thrombin or HRV 3C-cleavable C-terminal foldon tag with either a His_6X_ or SBP-His_8X_tag. All constructs were purified from supernatants by passage over Cobalt-TALON resin (Takara) followed by gel filtration chromatography on Superdex 200 Increase (GE Healthcare) in 10 mM Tris-HCl, 150 mM NaCl at pH 7.5. For BLI and crystallography the tags were removed using HRV 3C protease (ThermoScientific) and the protein repurified using Cobalt-TALON resin to remove the protease, tag and non-cleaved protein.

### Fab and IgG expression and purification

For Fab and IgG production the genes for the heavy- and light-chain variable domains were synthesized and codon optimized by Integrated DNA Technologies and subcloned into pVRC protein expression vectors containing human heavy- and light-chain constant domains, as previously described (^11,19^). Heavy-chain constructs for Fab production contained a non-cleavable His_6X_ tag; for IgG heavy constructs there was no cleavable purification tag. Constructs were confirmed by sequencing at the DNA Sequencing Core Facility at Dana Farber Cancer Institute. Fabs and IgGs were produced by transient transfection in suspension 293F or adherent HEK 293T cells using Lipofecatamine 2000 (Invitrogen) or polyethylenamine (PEI). Supernatants were harvested 4-5 days later, clarified by centrifugation. Fabs were purified using Cobalt-TALON resin (Takara) followed by gel filtration chromatography on Superdex 200 Increase (GE Healthcare) in 10 mM Tris-HCl, 150 mM NaCl at pH 7.5. IgGs were purified using Protein G Plus Agarose (ThermoFisher Scientific). Briefly, IgG supernatants were incubated overnight with agarose slurry, eluted with 0.1M glycine, pH 2.5 and normalized with 1M Tris-HCl, pH 8.0 and dialyzed against PBS buffer overnight.

### Crystallization and Data Collection

rsH4NBv1 HA1 head domain and K03.12 Fab were incubated at 1:1.5 molar ratio, respectively. The complex was isolated by size exclusion chromatography using a 24 mL Superdex Increase equilibrated in 10 mM Tris-HCl, 150 mM NaCl. Crystallization was achieved by hanging drop vapor diffusion at 18°C. Crystals were grown in 100 mM sodium citrate (pH 4.5), 20% (wt/vol) PEG 4000. Crystals were cryoprotected in mother liquor supplemented with 25% (vol/vol) glycerol and flash-frozen in liquid nitrogen. Data were collected at 0.999 Å with a rotation of 1° per image on the 8.2.2 beamline, Advanced Light Source, at Berkeley National Laboratory.

### Structure Determination and Analysis

The structure was determined by molecular replacement using PHASER ^26,27^ with the K03.12-A/Texas/50/2012 (H3N2)-head complex (PDB ID 5W08) as a search model (ref). Density-modified, NCS-averaged electron density maps were generated with DM (CCP4) and were used as guide for model building. Refinement of individual and group B factors was performed using PHENIX ^23^. Model building was done in COOT ^28^ and assessed with MolProbity ^29^. N-linked glycan stereochemistry was validated with Privateer ^30^. Figures were generated using PyMOL Molecular Graphics System (v1.8.0.0; Schrödinger LLC).

### Interferometry binding experiments

Interferometry experiments were performed using a BLItz instrument (forteBIO, Pall Corporation). Fab were immobilized on a Ni-NTA biosensor and cleaved rHA heads were titrated to obtain binding affinities. Initial, single-hit concentrations, were tested at 35µM for binding and then subsequent titrations for at least three different concentrations (chosen depending on the apparent *K*_*D*_ from the high concentration); the refined *K*_*D*_ was obtained through global fit of the titration curves by applying a 1:1 binding isotherm using vendor-supplied software. All experiments were performed in 10 mM Tris-HCl, 150 mM NaCl at pH 7.5 and at room temperature.

### ELISA

5-10ng of rHA FLsE were adhered to high-capacity binding, 96 well-plates (Corning) overnight in PBS. Plates were blocked with non-fat dried milk in PBS containing Tween-20 (PBS-T) for 1hr at room temperature (RT). Blocking solution was discarded and 10-fold dilutions of RBS-directed IgGs in PBS were added to wells and incubated for 1hr at RT. Plates were then washed three times with PBS-T. Secondary, anti-human IgG-HRP (Abcam), in PBS-T was added to each and incubated for 1hr at RT. Plates were then washed three times with PBS-T. Plates were developed using 1-Step ABTS substrate (ThermoFisher) and immediately read using a plate reader at 410nm. Data were plotted using Prisim 6 (GraphPad Software) and affinities determined.

### Mice

C57BL/6 mice were obtained from the Jackson Laboratory. JH6 mice were generated as described below. Mice were bred and maintained under specific pathogen-free conditions at the Duke University Animal Care Facility. Mouse immunization experiments were approved by the Duke University Institutional Animal Care and Use Committee.

### Generation of human J_H_6 murine model

In the human J_H_6 mouse model, human J_H_6 replaces mouse J_H_1-J_H_4, and human D2-2 replaces mouse DQ52 segment. The choice of human J_H_6 and D2-2 was based on the observation that a substantial fraction of human anti-RBS antibodies utilize these gene segments in CDR H3. To generate this mouse model, a knock-in construct, designed to integrate human J_H_6 and D2-2 into mouse DQ52-J_H_ locus as described above, was transfected into a mouse embryonic stem cell line that was derived from a F1 mouse (129Sv x C57BL/6). The construct integrated human J_H_6 and D2-2 segments into the IgH^b^ allele from C57BL/6 strain; the other IgH^a^ allele from 129Sv strain was unmodified. ES clones with correct integration were identified with Southern blotting. The ES clones with human J_H_6 and D2-2 were injected into mouse blastocysts to generate chimeric mice. Breeding of the chimeric mice gave germline transmission, and germline mice were used for the immunization experiments.

### Immunizations

J_H_6 mice (female, 8 to 12-week-old) were immunized with 20 μg of H1 SI-06 HAs in Alhydrogel® via footpad. Eight weeks later, cohorts of mice were boosted with 20 μg of either H1 SI-06 HAs or rsH4NBv3 HAs in Alhydrogel® via hock. Mice were sacrificed 8 days after prime or boost immunizations, respectively, and the draining, popliteal lymph nodes and sera were collected to measure GC and antibody responses, respectively.

### Luminex multiplex assay

Reactivity of mouse sera was determined by Luminex multiplex assay (cite Bajic and Maron Cell host microbe 2019) with modifications. Briefly, mixtures of antigen-conjugated (H1 SI-06 or rsH4NBv3 HAs) microspheres were incubated with serially diluted serum samples for 2 hours at room temperature or overnight at 4°C (for competition assay) with mild agitation. Samples were diluted in PBS containing 1% cow milk, 1% BSA, 0.05% Tween 20 and 0.05% NaN_3_. For competition, human monoclonal IgG1 CH67 (2 ng/ml) was added to the plates without washing and plates were incubated for 2 hours at room temperature with mild agitation. After washing, PE goat anti-mouse IgG or PE mouse anti-human IgG (both from Southern Biotech, the latter for competition assay) was added to the plates and incubated for 1 hour at room temperature with mild agitation. After washing, microspheres were resuspended in PBS containing 1% BSA, 0.05% Tween 20 and 0.05% NaN_3_ and fluorescent signals from each microsphere were measured in a Bio-Plex^®^ 3D machine (Bio-Rad).

## Data Availability

Coordinates and structure factors have been deposited in the Protein Data Bank under accession code PDB 6UR5 for the rsH4NBv1-K03.12 complex.

## Acknowledgements

We thank Stephen Harrison and members of his laboratory at Harvard Medical School for helpful discussions, in particular Kevin McCarthy. We thank Wenli Zhang, Dongmei Liao, and Xiaoe Liang at Duke University for technical assistance and mouse colony maintenance. BLI affinity measurements were carried out in the Center for Macromolecular Interactions at Harvard Medical School, directed by Kelly Arnett. We thank the beamline staff at the Advanced Light Source (Lawrence Berkeley Laboratory), beamline 8.2.2, for assistance in recording of X-ray diffraction data. We thank Yoana Dimitrova for help with x-ray data processing. Sequencing reactions were carried out with an ABI3730xl DNA analyzer at the DNA Resource Core of Dana-Farber/Harvard Cancer Center (funded in part by NCI Cancer Center support grant 2P30CA006516-48). This research was supported by NIH grants P01 AI089618 (to S.C. Harrison), U19 AI117892 (to G. Kelsoe) and R01 AI146779 (to A.G. Schmidt)

## Author Information

### Author Contributions

A.G.S. designed research; G.B., M.J.M., M,T., M.K. and A.G.S. performed research; G.B., G.K., M.K., and A.G.S. analyzed data; A.G.S. wrote the paper. G.B., M.J.M., M.T. G.K., and M.K. edited and commented on the paper.

### Competing financial interest

A.G.S., G.B. and M.J.M. have filed a patent application regarding the work published in this manuscript.

