## Supplementary Figures for "Structure-guided molecular grafting of a complex broadly neutralizing viral epitope"

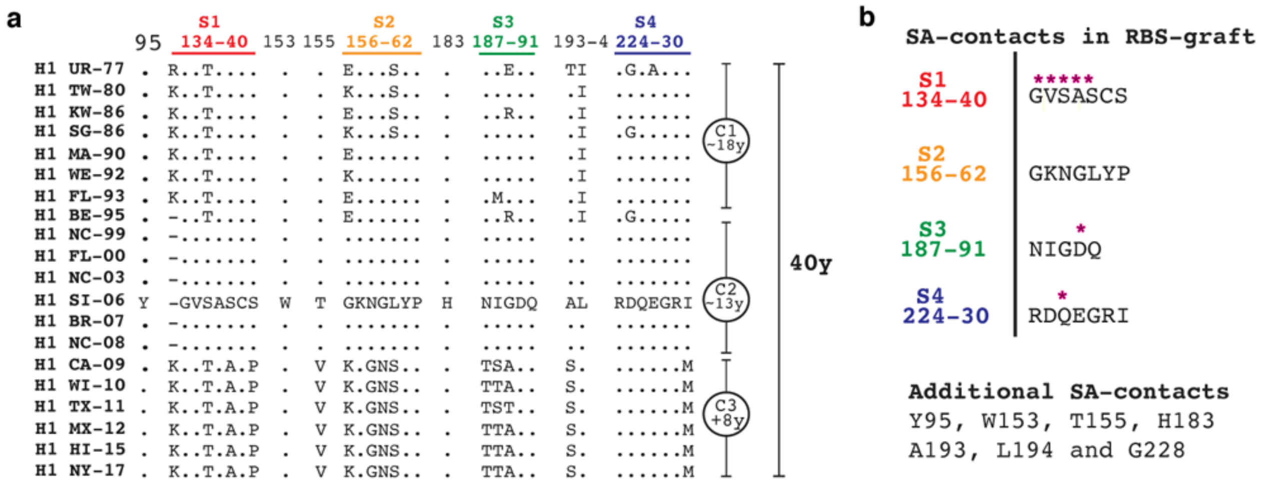


**Extended Data Figure 1: Conservation of the H1 HA RBS and critical SA contacts. a,** Sequence alignment of historical H1 RBS and critical residues comprising sialic acid (SA) contacts. The segments in the RBS graft are colored. Conserved residues (.) are in reference to H1 SI-06. The representative antigenic clusters (C1, C2, C3) of H1 isolates are listed with the numbering of circulating corresponding years (y). In order, the representative H1 isolates are: USSR-1977, Taiwan-1980, Singapore-1986, Massachusetts-1990, Wellington-1990, Florida-1993, Bejing-1999, Florida-1990, North Carolina-2003, Solomon Islands-2006, Brisbane-2007, North Carolina-2008, California-2009, Wisconsin-2010, Texas-2010, Mexico-2012, Hawai'i-2015, New York-2017. **b,** SA-contact residues in the RBS-graft segments are noted as asterisks (magenta) with the additional, conserved SA-contacts not present in the graft are listed.

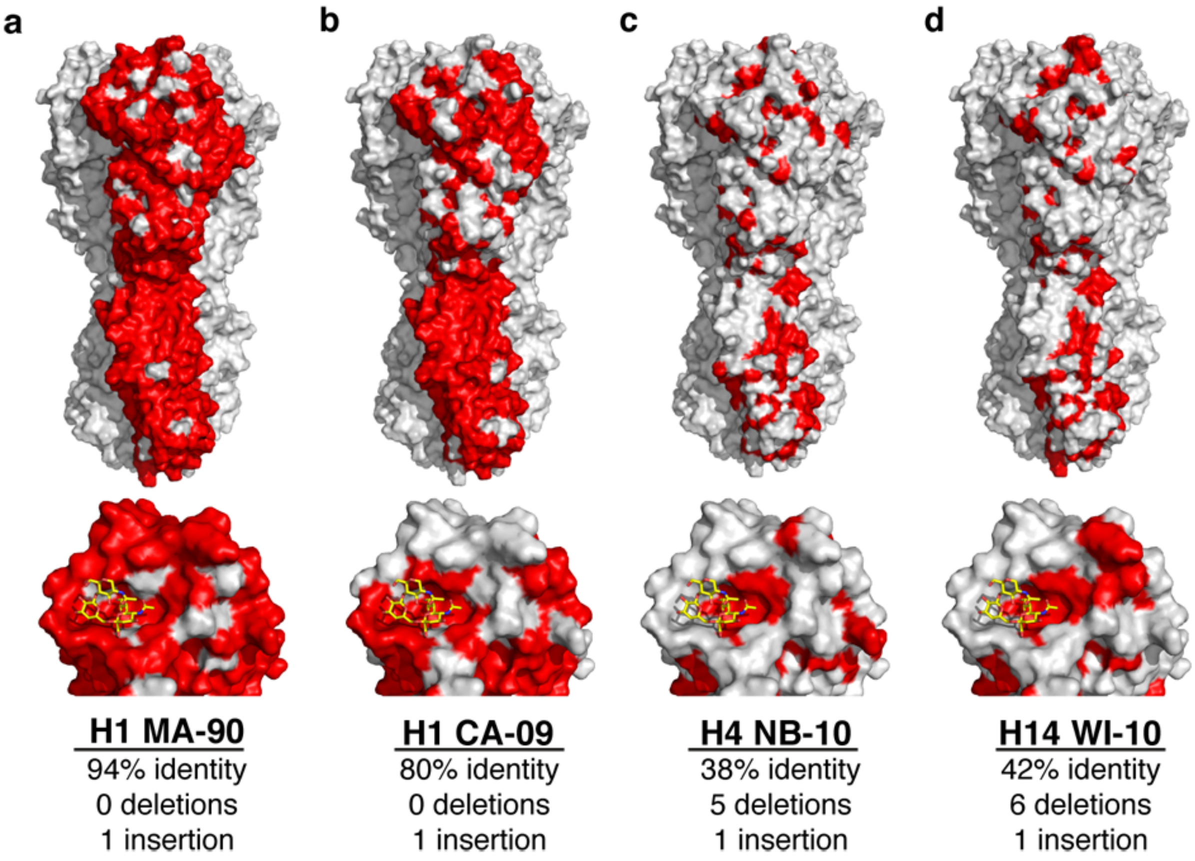


**Extended Data Figure 2: Conservation of HAs.** Using H1 SI-06 as reference, residue conversation is shown in red for historical H1 MA-90 and the new pandemic, H1 California/04/2009 (H1 CA-09) as well as the two acceptor scaffolds H4 NB-10 and H14 WI-10. Two views are shown: top is the HA trimer in spacefill with only one monomer colored red at points of conservation; the bottom is a close-up the RBS with a LStc molecule (stick-representation) docked for point of reference. The percent identity is in reference to H1-SI-06 with the total number of insertions and deletions listed. For both H1 MA-90 and H1 CA-09, the insertion is residue K133a.

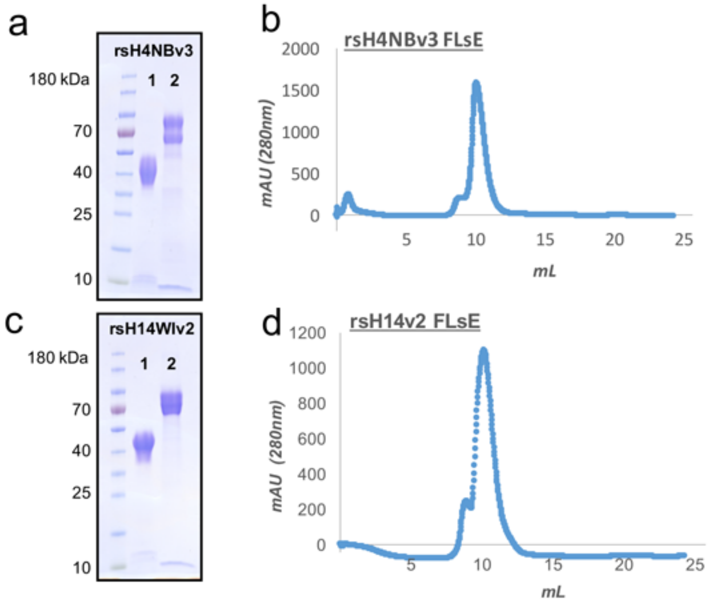


**Extended Data Figure 3: Biochemical characterization of the optimized rsHAs. a,** Coomassie-stained SDS-PAGE gel of the rsH4NBv3 head and FLsE constructs (marked "1" and "2", respectively). The doublet for the FLsE construct is proteolysis of the purification tags. A prestained protein ladder is in the first and the corresponding molecular weights (in kilodaltons) are marked. **b,** Representative FPLC trace using a Superdex 10/300 column of the FLsE construct. The trace monitors the absorbance (in mAU) at 280nm as a function of elution volume (mL). **c,** Coomassie-stained SDS-PAGE of the rsH14WIv2 constructs (labeled as in **a)**) and **d,** representative FPLC trace (similar to **d)**)

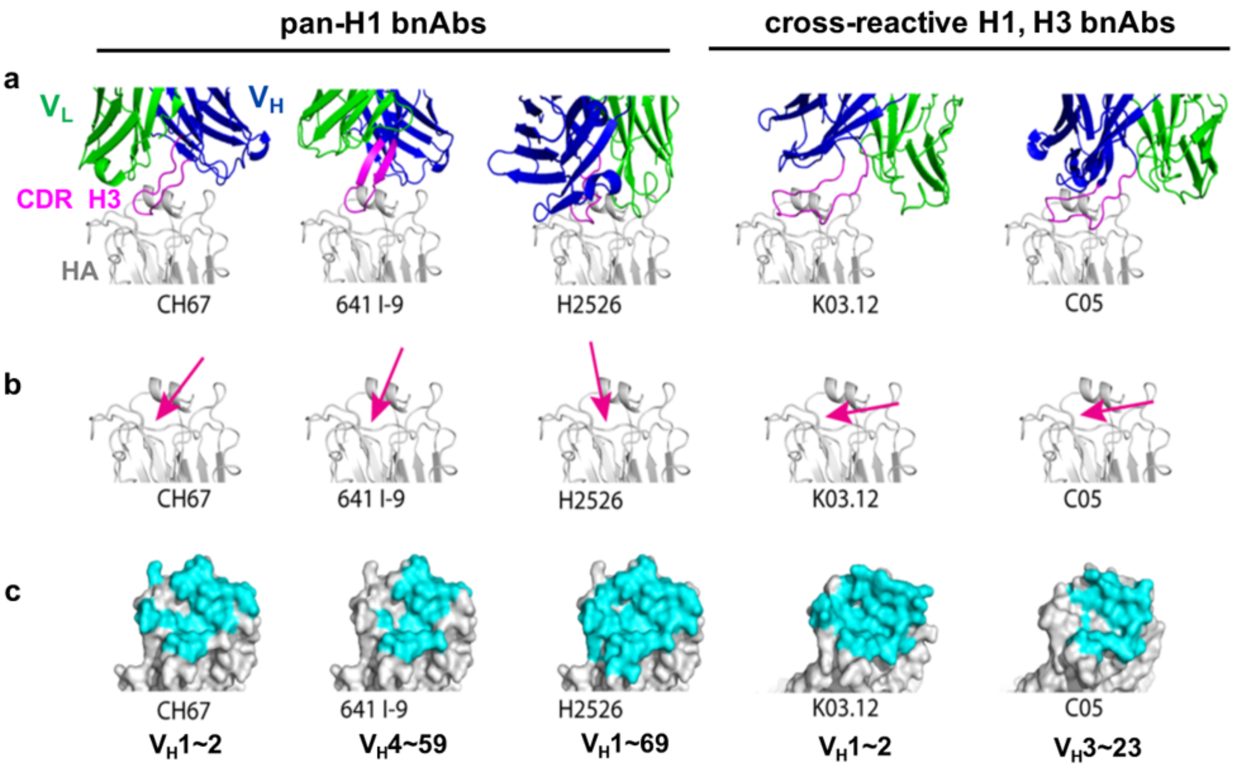


**Extended Data Figure 4: Antibody footprints of RBS-directed antibodies.** RBS-directed antibodies used in this study to obtain ELISA and BLI binding affinities. **a,** Crystal structures of CH67 (PDB 4HKX), 641 I-9 (PDB 4YK4), H2526 (PDB 4YJZ), K03.12 Fab (PDB 5W08) and C05 (PDB 4FQR) in complex with HA (silver). The variable heavy and light domains are colored blue and green, respectively with the CDR H3 in the antigen combining site shown in magenta. **b,** An approximate angle of approach of the CDR H3 of each antibody with the HA RBS. **c,** The overall footprint of each antibody with HA is shown in cyan with variable heavy ($V_H$) gene usage noted. All figures were created using PyMol.

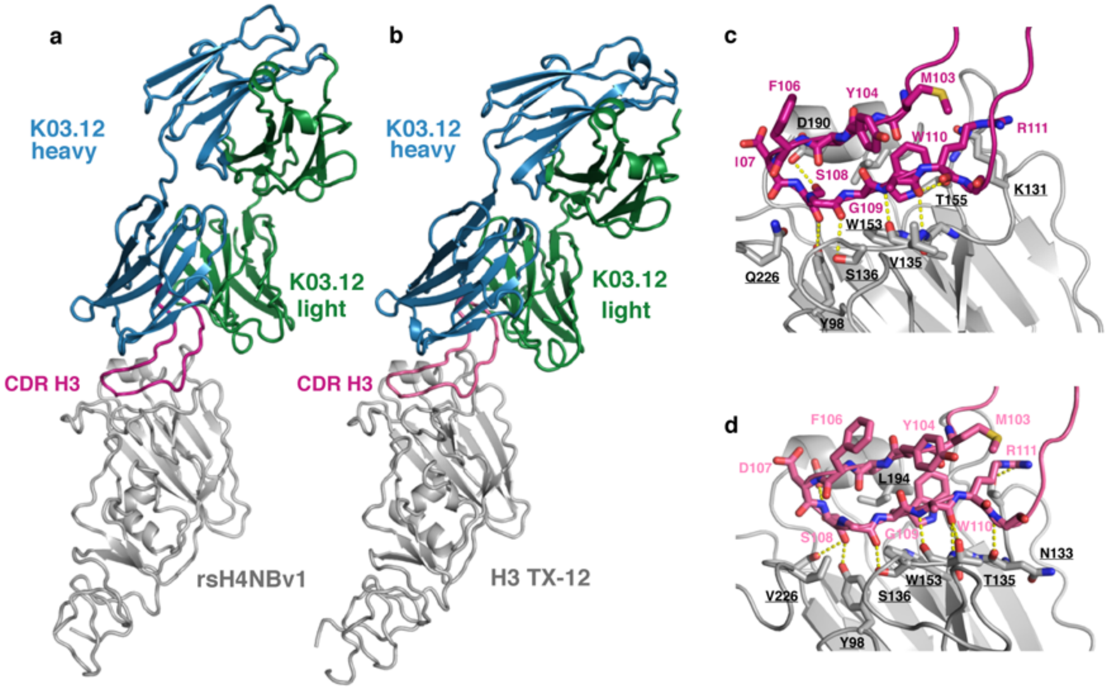


**Extended Data Figure 5: a,** K03.12 Fab (heavy and light chains are colored blue and green, respectively) in complex with rsH4NBv1 HA1 "head" (silver). The CDR H3 (magenta) is marked. **b,** K03.12 Fab (heavy and light chains are colored blue and green, respectively) in complex with H3 TX-12 (PDB 5W08) HA1 "head" (silver). The CDR H3 (magenta) is marked. **c, d,** Close-up of the antigen combining site. The CDR H3 (magenta) is shown in sticks with key interacting HA residues (silver). Hydrogen bonds are denoted in yellow, dashed-lines.

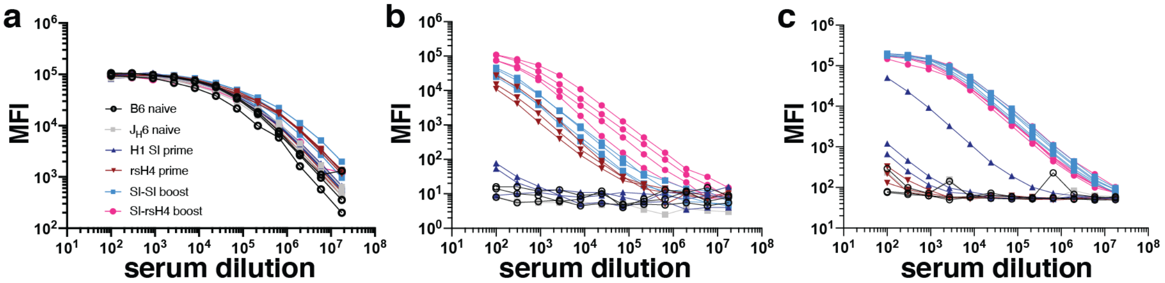


**Extended Data Figure 6: a,** Total IgG titer measurements of the different mouse cohorts. **b, c,** Serum reactivity measurements to rsH4NBv3 and H1 SI-06 HAs, respectively, using Luminex assay.

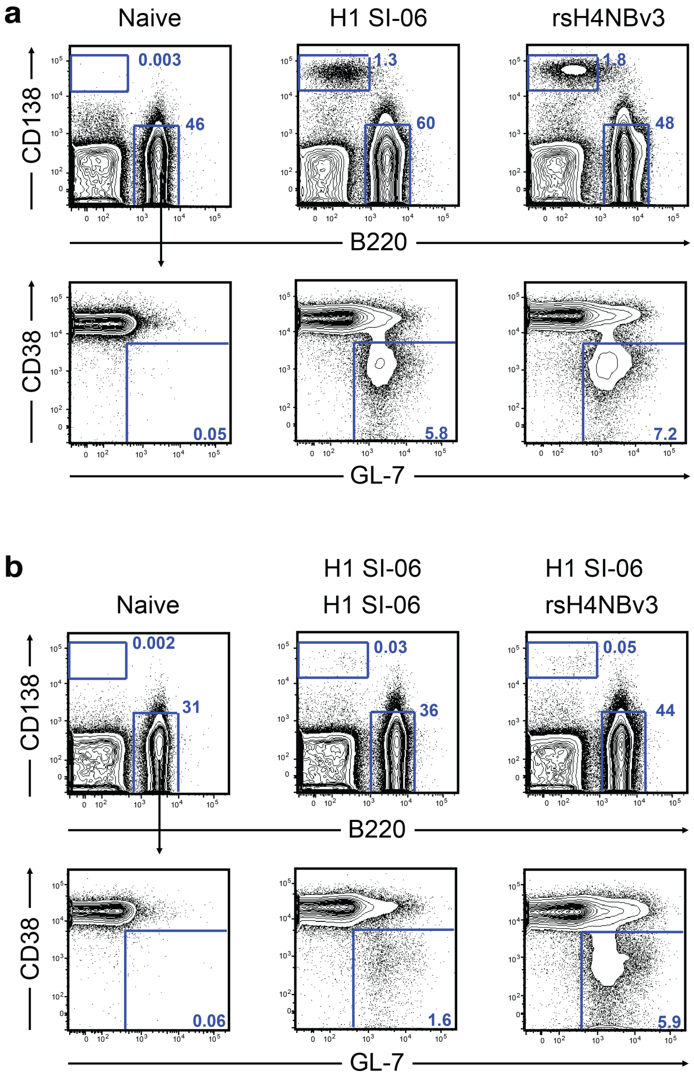


**Extended Data Figure 7: a,** Mice were immunized with recombinant H1 SI-06 or rsH4NBv3 HA in Alhydrogel® via footpad and the GC responses were assessed by flow cytometry 8 days after immunization. **b,** after initial immunization as in **a**, the mice were boosted with H1 SI-06 or rsH4NBv3 56 days later; the GC responses were assessed by flow cytometry 8 days after boost. Representative flow diagrams of B cells in the popliteal lymph nodes are shown.

| **Data collection and processing** | |
|---|---|
| Wavelength (Å) | 0.999 |
| Resolution range (Å) | 46.05 - 4.0 |
| Space group | C 1 2 1 |
| Unit cell a,b,c (Å)<br>α,β,γ (°) | 92.3, 141.5, 166.9<br>90, 102.6, 90 |
| Total reflections | 63073 (4479) |
| Unique reflections | 17320 (1493) |
| Multiplicity | 3.6 (3.0) |
| Completeness (%) | 97.5 (85.8) |
| Mean I/sigma(I) | 4.51 (1.52) |
| $R_{merge}$ | 0.29 (0.73) |
| $R_{meas}$ | 0.34 (0.89) |
| $R_{pim}$ | 0.17 (0.49) |
| CC1/2 | 0.95 (0.54) |
| CC* | 0.99 (0.84) |
| **Refinement** | |
| $R_{work}/R_{free}$ | 0.29/0.32 |
| RMS bonds (Å)/angles(°) | 0.003/0.70 |
| Ramachandran favored/outliers (%) | 94.65/0.14 |
| Average B-factor | 97 |

Statistics for the highest-resolution shell are shown in parentheses.

**Table S1 Crystallographic Data Collection and Model Refinement statistics.**
